# Revealing brain network dynamics during the emotional state of suspense using topological data analysis

**DOI:** 10.1101/2024.01.29.577820

**Authors:** Astrid A. Olave, Jose A. Perea, Francisco Gómez

## Abstract

Suspense is an affective state ubiquitous in human life, from art to quotidian events. However, little is known about the behavior of large-scale brain networks during suspenseful experiences. To address this question, we examined the continuous brain responses of participants watching a suspenseful movie, along with reported levels of suspense from an independent set of viewers. We employ sliding window analysis and Pearson correlation to measure functional connectivity states over time. Then, we use Mapper, a topological data analysis tool, to obtain a graphical representation that captures the dynamical transitions of the brain across states; this representation enables the anchoring of the topological characteristics of the combinatorial object with the measured suspense. Our analysis revealed changes in functional connectivity within and between the salience, fronto-parietal, and default networks associated with suspense. In particular, the functional connectivity between the salience and fronto-parietal networks increased with the level of suspense. In contrast, the connections of both networks with the default network decreased. Together, our findings reveal specific dynamical changes in functional connectivity at the network level associated with variation in suspense, and suggest topological data analysis as a potentially powerful tool for studying dynamic brain networks.

## 1 Introduction

Suspense is an affective state associated with conflict, dissonance, instability, or uncertainty regarding an emotionally significant circumstance. On some level, the particular event is not susceptible to influence or control, which motivates future-oriented expectation or prediction and desire for a resolution [1]. Researchers have identified subcortical regions and some regions from large-scale networks: the salience, fronto-parietal and default networks to be involved in the experience of suspense [2–4]. Recent evidence confirms that these networks and the ventral-attention network characterized five different activation brain states that occur during suspenseful film viewing with a likelihood of state’s appearance near suspense peaks. [5]. However, activation and deactivation may be insufficient to understand the intrinsic neuronal mechanisms of suspenseful experiences, as evidenced by the recently described interactions of large-scale networks underlying the human emotional response [6–9]. In particular, changes in the network-level interactions between salience, frontoparietal, and default networks are related with the presence of stress or anxious anticipation [10–12], emotional states akin to suspense. Interestingly, psychological models of suspense anticipate this kind of interaction [1], relating the emotional brain processing to network-based dynamics.

fMRI functional connectivity (FC), understood as a description of synchronizations among neurophysiological time series of different regions, may help to characterize brain networks involved in suspense processing. Commonly, FC analyses rely on a *static brain network* representation which contains measures of pairwise interactions between regions. In this approach, each interaction corresponds to the correlation between the full time series produced by each brain area. This FC representation provides a “average” synchronization over the entire temporal span, which may obscure possible changes in the brain dynamics occurring on shorter time scales of the order of seconds or even milliseconds, which have proved to be relevant for cognitive processing [13].

Recently, FC information has been complemented with dynamic functional connectivity (dFC) analyses; the latter describe how brain interactions may change over time. Most of these approaches quantify pairwise interactions among brain regions in short time windows and then compute the dynamics of change of these interactions over time. Different methods provide these dynamical FC descriptions, including sliding windows, time-frequency analyses, dynamic graphs, point-processes, and co-activation patterns, among others [14, 15]. Nevertheless, most of these works focus only on describing a set of dynamical states without considering possible transitions among them. An approach that can be insufficient to unveiling the high-order dynamical organization of the brain, expected during the ongoing emotional processing induced by suspense experiences [1].

Topological data analysis (TDA) is a field emerging from convergent works in applied (algebraic) topology and computational geometry. TDA offers a variety of methods to robustly infer qualitative and quantitative information about the structure of data, both at the local and global scales. One of its approaches, Mapper [16], has been recently used to reveal the organization of complex brain dynamics during resting-state [17] and ongoing sensory and cognitive activity processing [18–20]. Mapper reduces a high-dimensional dataset, like one describing different brain states, into a simplicial complex - a high-dimensional analog of a graph [21]. In contrast to other methods, Mapper does not collapse data in space of time, does not require a prior knowledge of the number states nor a strict characterization, is under small deformations, thus, is more robust to noise, it is stable trough different scales of resolution and its graphical output allows to distinguish the relations between the data, for example, the emergence and transitions between brain states. As a result, TDA-based methods have been useful in describing fMRI brain dynamic organization. Nevertheless, to our knowledge, there is no topologically-related description of the underlying dynamics exhibited by FC during suspense processing.

Our present work examines the changes in functional connectivity between large-scale networks over time in the presence of emotional experience of suspense using topological descriptors. In contrast to previous studies focused on the search of regions involved in suspense processing [2–4] or in the description of brain states emerging from region activation [5], we characterize the brain states trough FC dynamics of the salience, fronto-parietal, and default mode networks, given their involvement in suspense and anxiety processing. Then, we use Mapper to encapsulate the FC dynamic’s underlying topology by a graph which provides a high-order description of the suspense brain’s dynamic. From Mapper’s representation we distinguish key topological features like relevant states engaged in the dynamics, transitions between them, and groups of dynamical states or communities.

The topological features were then related to the suspense level reported by independent film watchers. Reliability and validation analysis were also considered to study the robustness of the results [22]. The joint analysis of the level of suspense and the topological description of the brain dynamic suggest a correlation between the variations in the intensity of suspense with changes in the functional connectivity within and between salience, fronto-parietal, and default networks, emerging on the topological representation. In particular, the FC between the salience and fronto-parietal networks increased with the level of suspense. In contrast, the connections of both networks with the default network decreased. Altogether, results demonstrate that the emotional state of suspense is associated with dynamic FC interactions between large-scale networks, which may be unveiled by topological-based approaches, allowing to describe a complex brain dynamic related to emotional processing.

## 2 Results

### Revealing dynamic functional connectivity using topology

We analyzed an open dataset acquired by the Cambridge Center for Aging and Neuroscience (Cam-CAN) [23, 24] and processed by Schmälzle & Grall [4]. The fMRI data was collected while participants watched an edited version of Alfred Hitchcock’s “Bang! You’re Dead”, a black and white television drama portraying a little boy who plays around with a charged revolver, thinking it is a toy gun (Fig. 1**A**). In addition, a different set of participants watched the same clip and “continuously evaluated the degree of suspense” they were experiencing. The average degree of reported suspense is provided by Schmälzle & Grall [4] (Fig. 2**B**).

**Figure 1.**
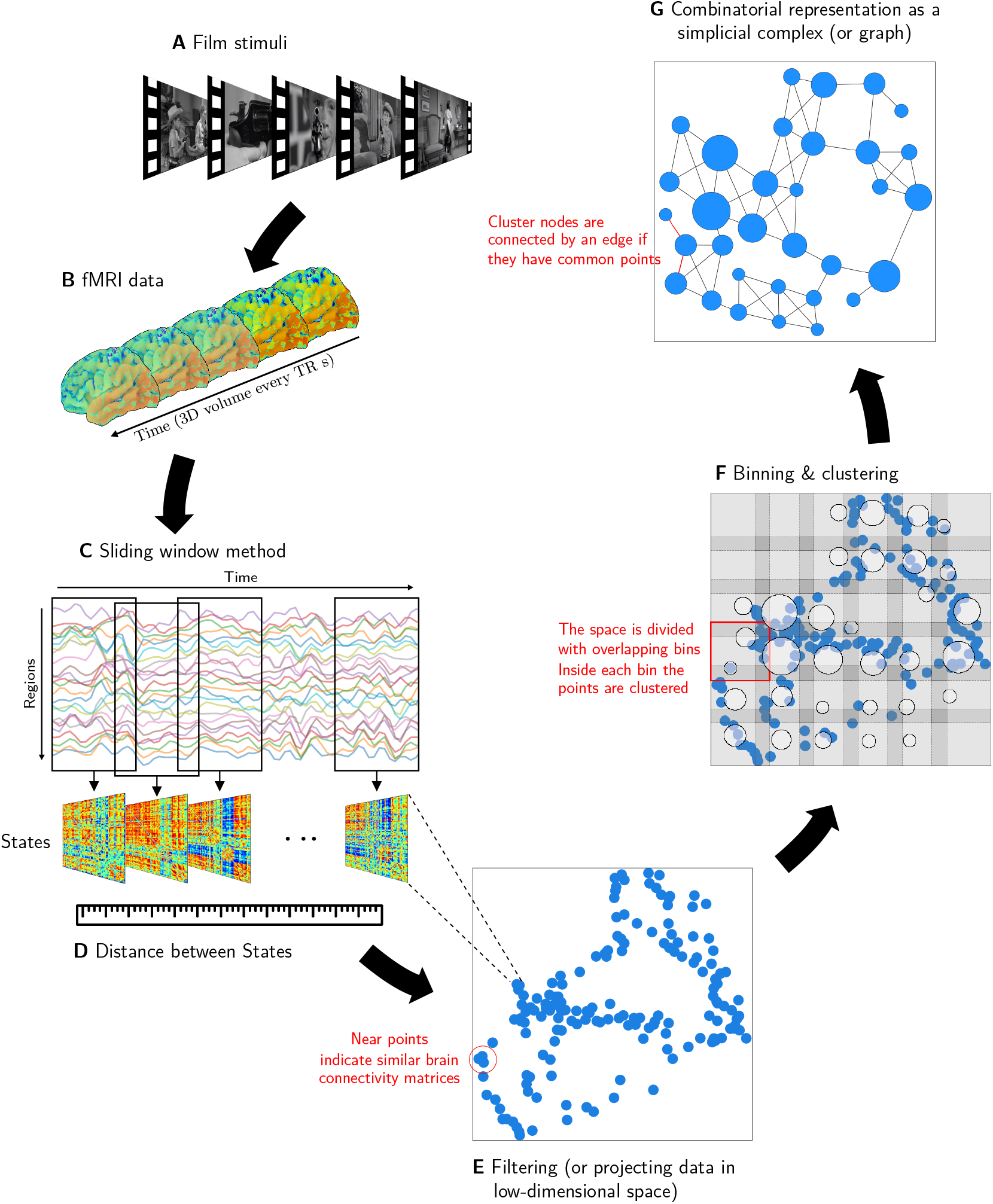
The schematic flowchart of analysis applied to fMRI data. **A** Participants watch a film while being scanned **B** The fMRI dataset is pre-processed and represented as time-series. **C** The SW is applied to compute the functional connectivity over time. **D** A matrix of similarities between connectivity matrices (states) is calculated. `Mapper algorithm`: **E** The set of states are projected into a two-dimensional space using the similarity matrix. **F** The space is divided into smaller bins determined by the number in each dimension and the percent of overlap between them. Next, partial clustering is applied within each bin. **G** To create a compressed combinatorial representation (graph) each cluster is treated as a node and two nodes are connected by an edge if they share data points.

**Figure 2.**
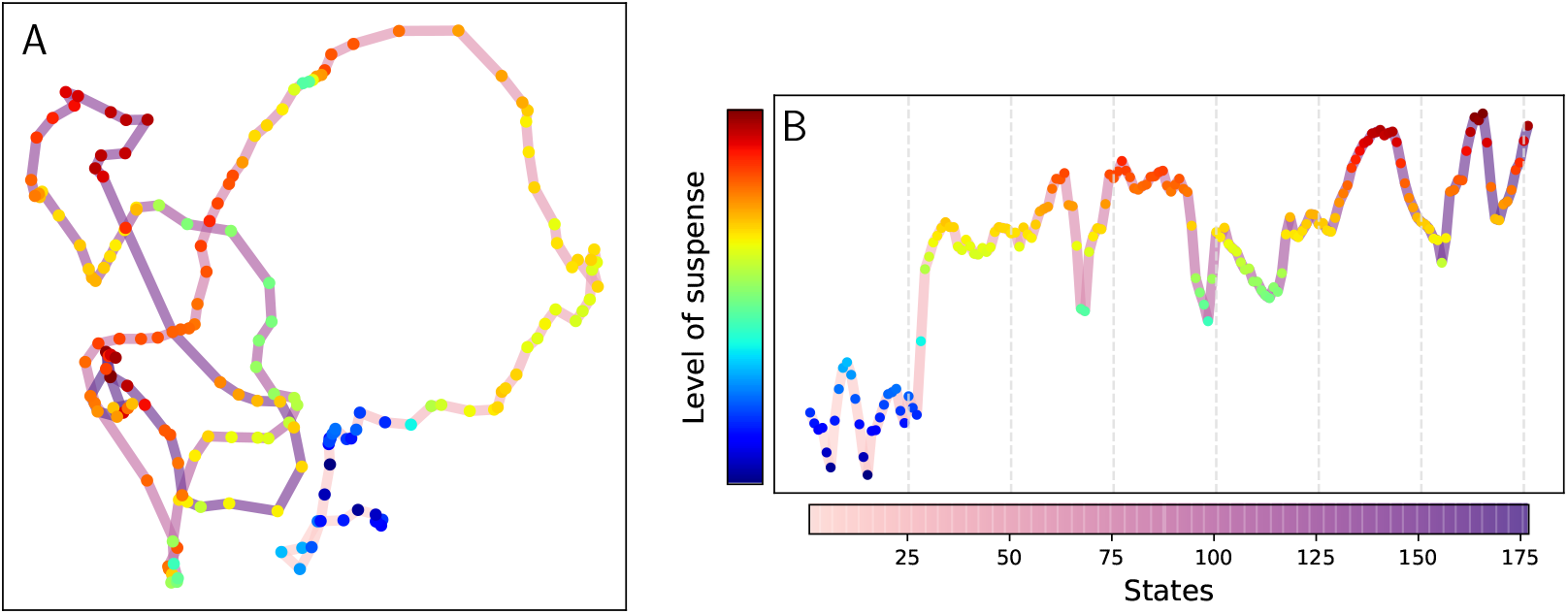
Panel **A** Isomap filtering. The colored dFC states are joined by a degraded pink-purple line representing the flow of time, i.e., state *i* is joined with a segment with state *i* − 1 and a slightly darker segment with state *i* + 1. Panel **B** reported level of suspense for the states.

Each participant’s fMRI data was represented by a matrix where the columns corresponded to time frames (or volumes), and rows to the activity magnitudes of regions from the salience, fronto-parietal, and default networks. Then, we characterized the dFC using the *sliding window method* (SW) (Fig. 1**C**). This procedure started by calculating the FC of the regions between volumes 1 and *W* (window size), generating a connectivity matrix that describes the FC *state* exhibited by the brain during the corresponding temporal frame. Then, the window was shifted (slid) by a step *S* to repeat the calculations between *S* and *W* + *S*. The process was then repeated until the window covers all the volumes. Thus, a set of connectivity matrices represented the temporal evolution of the whole brain FC. We used the SW over all individuals and computed the average of all connectivity matrices for each state.

A key idea of TDA is that the notion of similarity between the data points (distance) should be the starting point to reveal the shape of the data [25]. Hence, the first step for the topological description of the dynamic was to construct a metric space, where the data points corresponded to the connectivity matrices, and the dissimilarity captures the distinctness between FC states (Fig. 1**D**). The Manhattan distance, which is the sum of the absolute differences in the functional connectivity for each pair of regions, was selected to compute this dissimilarity. This metric was selected because it promotes the sparsity thereby mitigating the course of dimensionality naturally expected when comparing the high-dimensional data associated with the connectivity matrices [26].

Afterward, Mapper was used considering the data space of the matrices representing dFC. More specifically, 1) filtering, 2) covering with overlapping bins, 3) clustering, and 4) creating the graph (Fig. 1**E-G**).

The data points were filtered, i.e., they were mapped in a low-dimensional space (filter space) using a function (filter). The filter was chosen such that the time points with similar FC states in the original dataset were mapped “close” in the filter space. Meanwhile, different points ended up “far” from each other (Fig. 1**E**). *Isometric Mapping* (Isomap), a non-linear dimensionality reduction method is used to performed this reduction. The method is informed by the distance in the metric space and using geodesic distance to learn the global geometry of the data set and converge asymptotically to the true data structure [27]. This filter aimed to preserve and highlight the neighboring states exploiting the local linearity of the dataset. The filtering result is shown in Figure 2**A**, where each dot represents a dFC state. For a qualitative examination, the states were colored, indicating their level of suspense, and the dots were joined with a line that marks the flow of time. Figure 2**B** shows the level of suspense for the states as a guide to better recognize the corresponding state in the filtering.

Subsequently, the filter space resulting from Isomap was divided into pieces that covered the collection of points. Each piece from the covering was decomposed into groups using a clustering algorithm that agglomerates data points using the *original Manhattan metric*, i.e., each piece corresponded to clusters of similar points. It is worth noting that this clustering step was calculated on the high-dimensional space. Thus, it was possible to recover the structural information missed after filtering into a low-dimensional space. Hence, the filter space allows us to slice the data into meaningful, though smaller pieces, and compress the data instead of trying to apply a clustering approach over the whole set. For the clustering, the single-link agglomerative hierarchical method (Eq. 4) was used because it is not restricted to data in Euclidean space and does not require specifying the number of clusters beforehand (Fig. 1**F**).

Theoretically speaking, Mapper does not restrict the shape of the pieces, but regular coverings are preferred, for example, rectangles, hexagons, or balls [16, 28]. The proposed analysis relied on *rectangles* uniformly distributed in the parameter space. The covering has two parameters: the number of intervals (*n*), in which each dimension of the parameter space is divided, and the percentage of overlap between the intervals, called overlap (*o*). These parameters determined the coarseness of the future Mapper output: low values of *n* would generate a small number of nodes, each containing probably a large amount of data points. Thus, there is a risk of losing structural information. On the other hand, high values of *n* result in a computationally expensive graph since it would generate nodes that do not carry additional information about the underlying structure. Meanwhile, high values of *o* could result in spurious connections (edges) on the combinatorial object, and low values of *o* could lose important connections, again losing structural information.

As the choice of different parameters may lead to different and even misleading outputs, both parameters were tuned to replicate the most persistent homology features of the metric space of the data (Fig. 1**D**) before the filtering step. In simple words, the 0-homology, *H*_0_, corresponds to connected components, and the 1-homology, *H*_1_, to 1-dimensional holes, i.e., cycles or circles, emerging in the data space. For this purpose, another topological tool called *persistent homology* was used to determine the topological features, namely, the elements of *H*_0_ and *H*_1_, that most persisted along different scale levels. Then, the Mapper parameters were chosen to reproduce these features. The details of this analysis are elsewhere presented [21].

Briefly, for a given real number *ϵ*, we associate a simplicial complex derived from the data points and we compute its homology. Subsequently, we repeat this algorithm for 0 *< ϵ < R*. The homological classes that *persist* through a large range of *ϵ* are considered to reveal important homological aspects of the dataset.

The results are presented in the persistence diagram (Fig. 3). In the figure, each dot represents a homological class, for example, each blue dot is a connected component. The *x* coordinate of the dot corresponds to the *ϵ* value when the homology class appears (birth) and the *y* the *ϵ* value when the homology class vanishes (death). The horizontal dashed line represents *R*. Thus, the farther the dot is from the diagonal, the more persistent is. We have encircled the most persistent classes in the Figure 3. Let us call *persistence* the difference between death and birth. With this term, we have one connected component with *R* persistence and one circle with persistence 97 (from 125 to 222) with the rest of circles not surpassing 25 in persistence. Therefore, mathematically speaking, we could infer that over a large range of scales there is a circle-shaped aspect to the data, since #*H*_0_ = 1 and #*H*_1_ = 1, *is the homology of a circle*.

**Figure 3.**
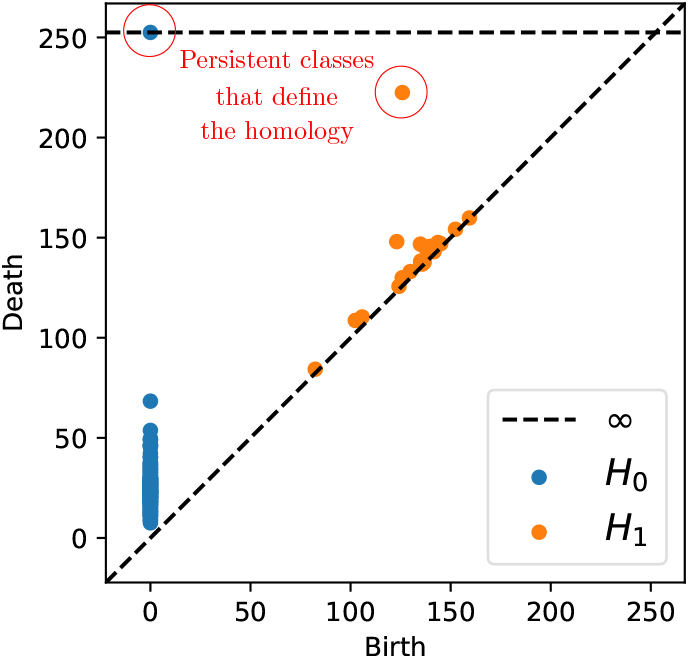
Persistence diagram presenting the homology of the set of States. *H*_*i*_ stands for *i*-homology.

Finally, to form the Mapper final graph, the former local clusters were considered as vertices, and edges connecting two vertices were included if they have points in common (Fig. 1**F**). Note that clusters can indeed share time points as the bins overlap.

Figure 4 shows the final graph representing the dFC. This graph possesses 52 nodes (as opposed to 176 states) and 100 edges. For qualitative analysis, the size and color of the nodes are considered in the vertices based on the number of states they contain. In this way, it is possible to observe the flow of time **A** and its relation with the suspense **B**.

**Figure 4.**
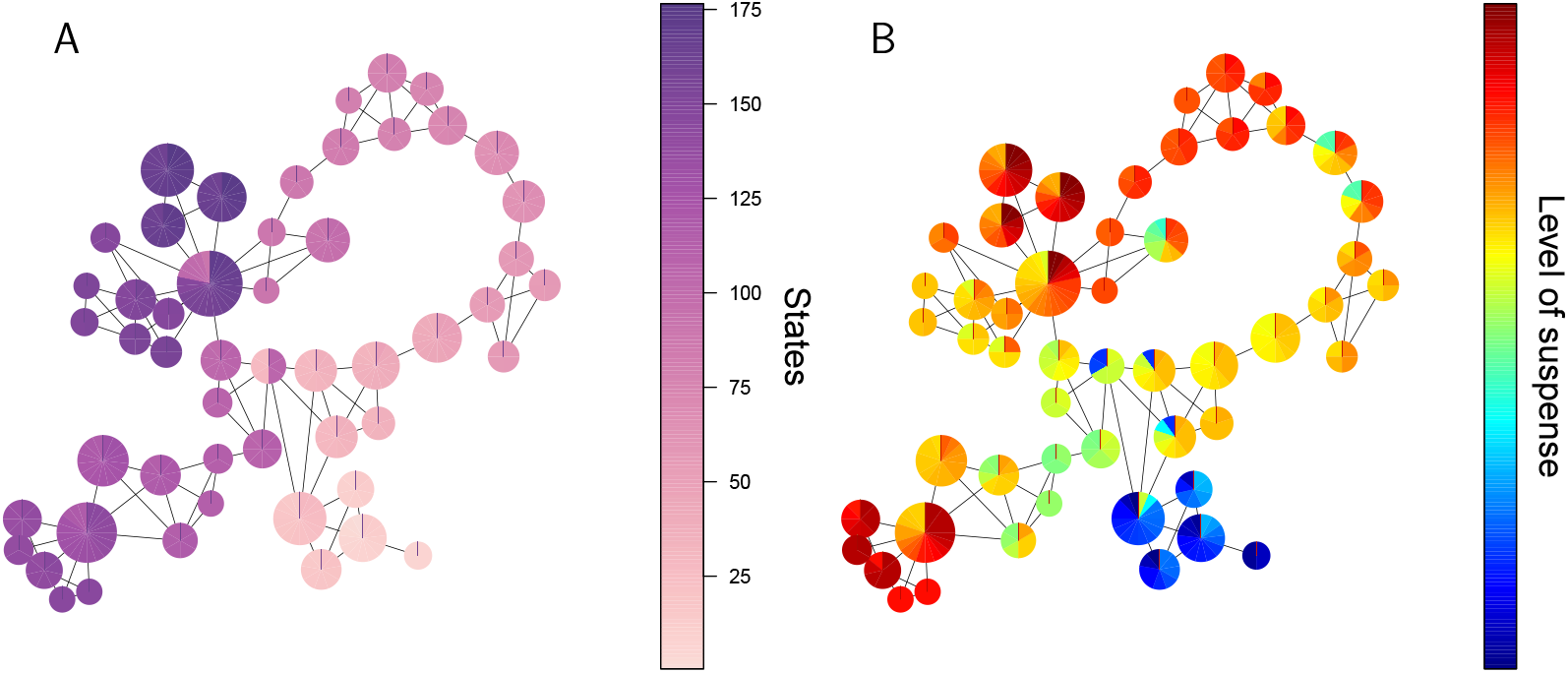
Final `Mapper graph.` Each node is sized in proportion to the number of contained states. The nodes are colored using a pie chart to indicate the range of **A** the index and **B** the levels of suspense among the states of the node.

As observed, the Mapper method maps the most similar states into a common node, while the closeness to other elements is represented by the edges between the nodes. This graph provides rich and robust representation of how the brain dynamically transitions across different FC states throughout the experience of suspense.

### Structural organization of suspense processing

Once the dFC is represented as a graph, it is possible to provide insights into the particular structural organization of the complex dynamic of the suspense processing. For this, we investigated the emergence of communities in the graph. A group of nodes is called a community when these nodes present greater connectivity among group members compared to the connectivity with nodes in other communities [29].

Figure 5 shows the communities found in the Mapper graph. These communities were determined using random walks [29]. Intuitively, random walks on the graph tend to get “stuck” in communities as they are densely connected, featuring the nodes that belong to the same community. Additionally, the graph allows us to visualize the temporal evolution of the brain from one community to another. For example, it highlights the *bridge* nodes, nodes connecting one group to the next. Moreover, it is possible to identify the FC states that are responsible for the connection between the bridge nodes, called *transition* states. For instance, nodes 26 and 37 connect the blue and orange communities because they share three states. Hence, those three FC states are transition states.

**Figure 5.**
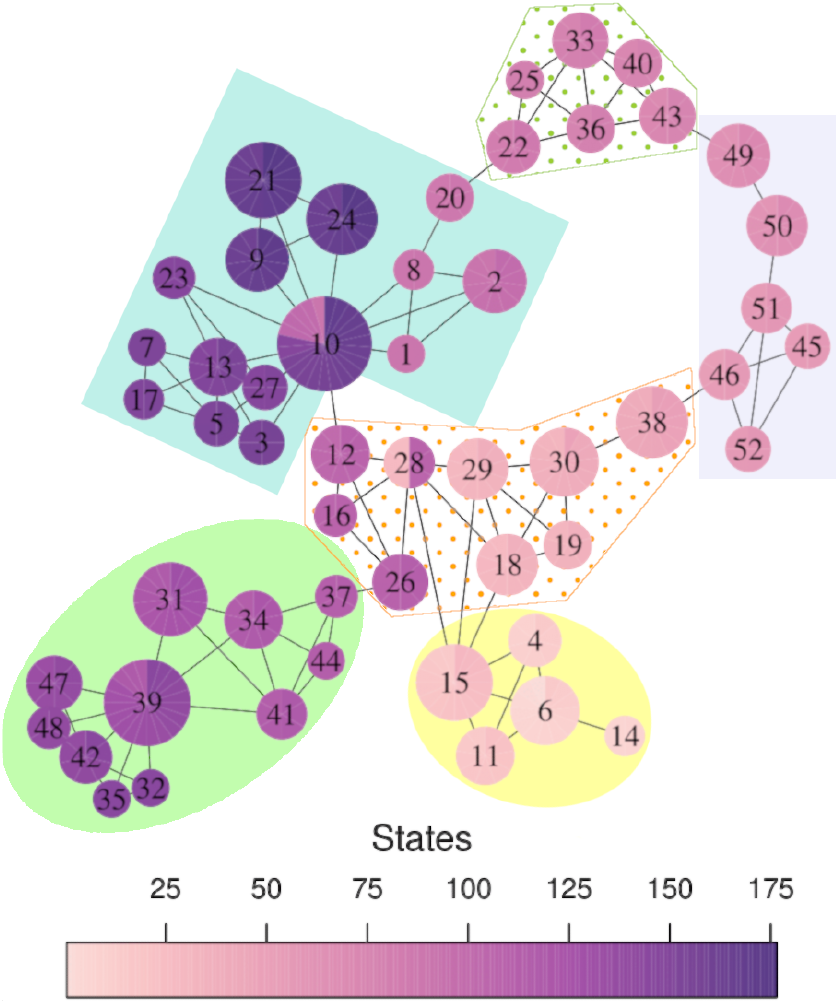
`Mapper` graph. The nodes are numbered and colored, denoting the indices of States. In different colors are enclosed the communities found in the graph

Furthermore, most of the bridge nodes are connector nodes (nodes that, after being removed, change the number of connected components) or cyclic nodes (nodes that, after being removed, change the number of cycles), which are key for preserving the shape homologically equivalent to a circle.

Thus, transition states inside connector nodes are the unique way the brain could enter or leave these communities. Meanwhile, states inside cyclic nodes allow navigation through these groups in two different directions (without considering the time constraint). In particular, states inside 10 and 28 are transition states that “jump” from phases that are temporally disconnected.

In addition, the quality of the community structure was computed using the function of modularity *Q* (Eq. 6), resulting in a value of 0.72, which indicates a strong community structure. Moreover, validation against the null model shows that this community organization present in the Mapper graph arises from the dynamic variation of the FC in the fMRI scans.

### Linking the graph to dynamic functional connectivity

With the purpose of linking the graph properties with the dynamic connections between the large-scale networks, the FC trough time within and between all pairs of large-scale networks was computed using connectivity weights, the average of the FC of the regions in each network. For this, the average connection between the regions in each network was considered, as observed in Eq 7. Specifically, six-time series corresponding to three within-networks weights (salience, fronto-parietal, and default) and three between-network weights (salience & fronto-parietal, salience & default, and frontoparietal & default) were computed. This analysis measures how strong the connections within and between networks were for each state.

For qualitative analysis, each vertex of the Mapper graph is colored using a pie char to indicate the range of connectivity weights within- and between-networks among the states of the node Fig 6. Remarkably, the communities are related to the level of connections between the networks. For instance, in the green solid circle community the level of connections between the default network and the salience and fronto-parietal networks are weak. Meanwhile, the doted green trapezoid community has states with a strong connection between the salience and fronto-parietal networks and within the fronto-parietal network. At the same time, the remaining interactions have a low level of connection.

**Figure 6.**
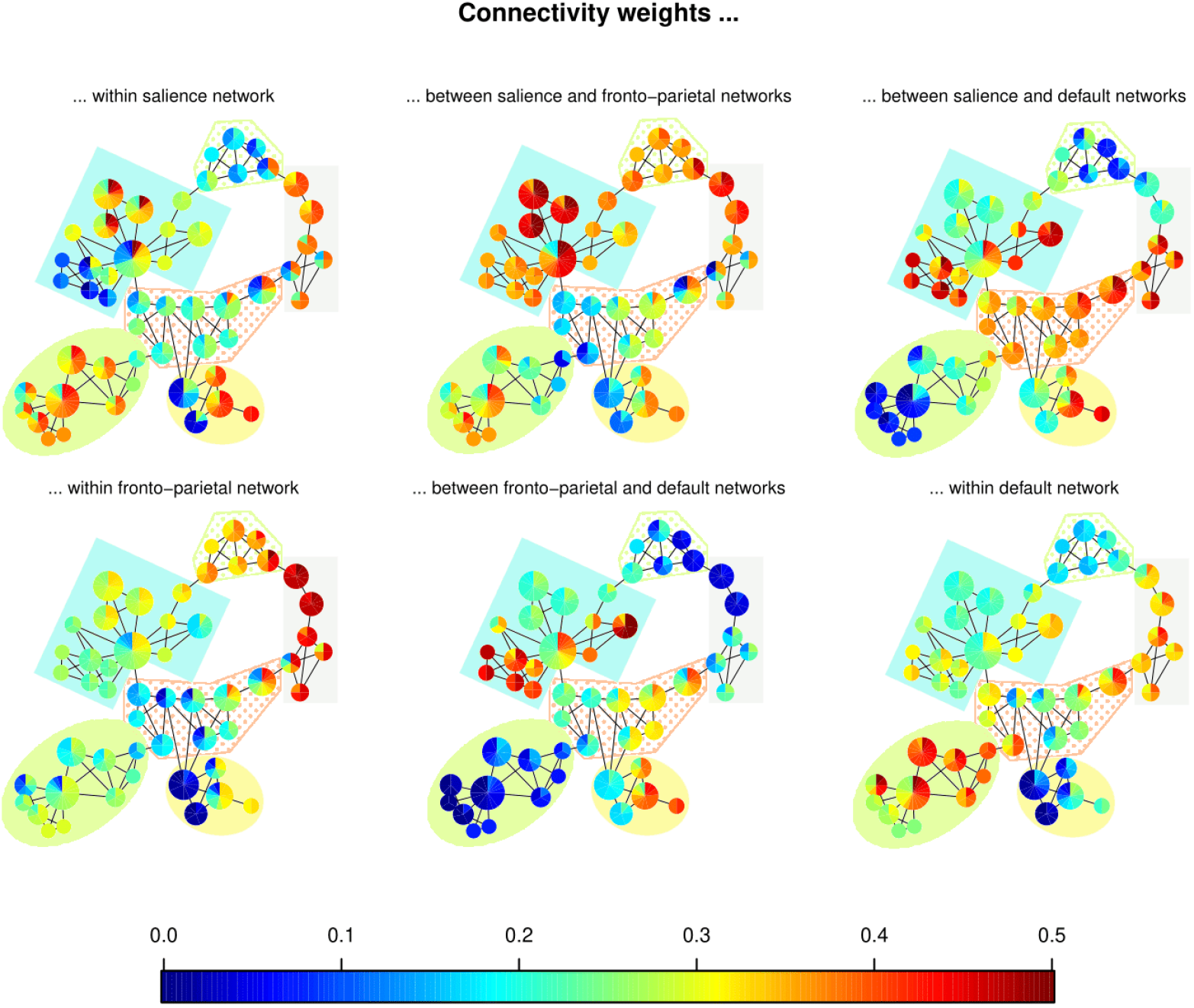
Mapper graphs. The nodes are colored using a pie chart to indicate the range of connections’ weights within and between networks. The communities found in the graph in 5 are enclosed in different colors.

Along with this analysis, the states were grouped by applying a complete-link hierarchical clustering (Eq. 5) using the geodesic distance between them. For this, a data-driven method was used to recognize similar groups but detected a single cluster. Hence, we manually chose the cut-off height to obtain the three clusters shown in Table 1.

**Table 1:** Table of intervals of States forming the groups clustering directly from the distance matrix.

| Cluster 1 | Cluster 2 | Cluster 3 |
| --- | --- | --- |
| 1-31 | 32-59 | 60-79 |
| 80-114 |  | 115-145 |
| 146-176 |  |  |

As expected, there is a correspondence between the intervals defined by the communities and those in the clusters. The intervals defined by the clusters were colored together with the intervals found in the communities of the
graph. In addition, orange and blue communities highlight how the bridge states inside nodes 10 and 28 connect the
intervals in cluster 1.

In a similar way, each time series of connectivity weights was associated with the communities to which it belongs (see Fig. 8). Notably, the intervals defined by the communities are also related to trends in the connections. For instance, in the lavender interval (52-72), the strength of all connections involving the default network decreased. Meanwhile, the other three pairs of networks showed an increasing behavior. Similarly, the limit between the green and blue intervals (145/146) corresponds the sudden increase in the connections between the default and the other two networks.

**Figure 7.**
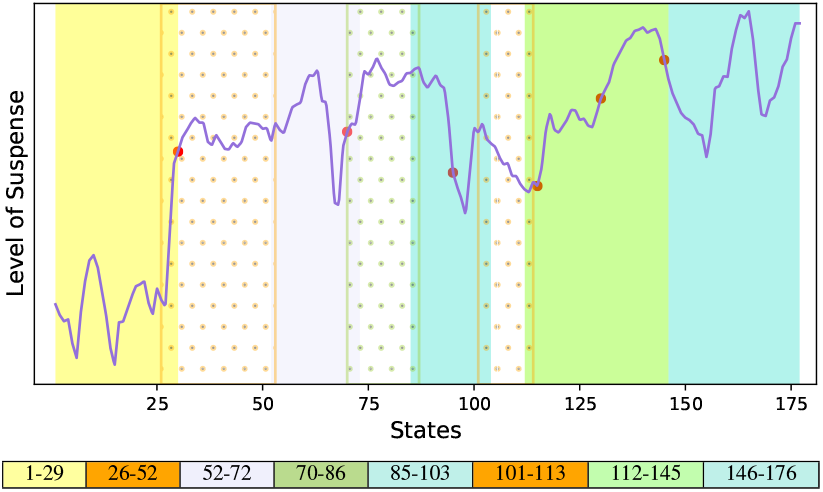
Level of suspense of the states. The vertical colored rectangles indicate the intervals associated with the communities in the `Mapper` graph. The red dots are change points of the time series.

**Figure 8.**
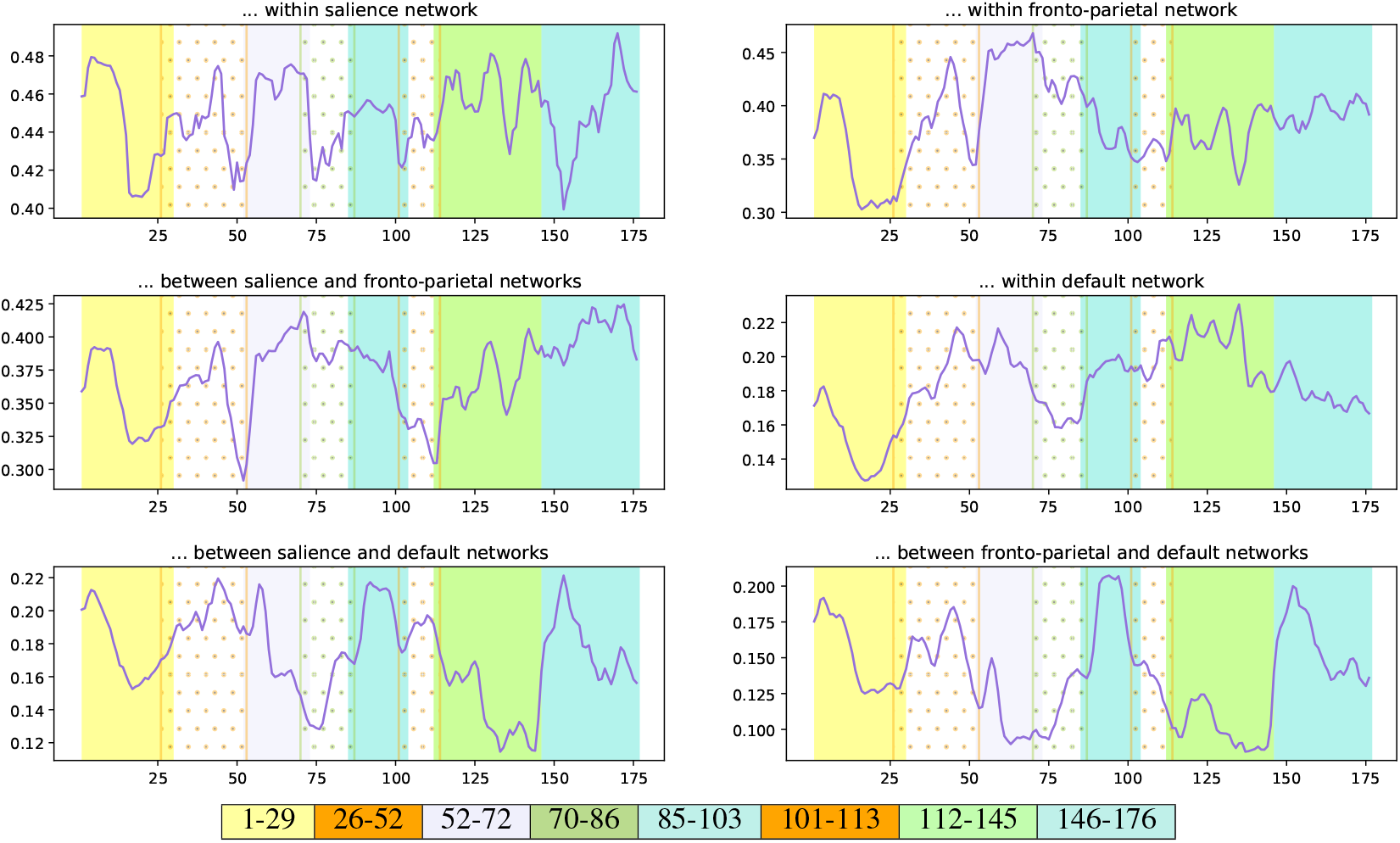
Time series of connections’ weights within- and between-networks (purple) and the average value (dashed gray). The vertical colored rectangles indicate the intervals associated with the groups of states from the `Mapper` graph in Figure 5.

### Associating dynamic functional connectivity with suspense

To test the hypothesis that dynamic functional connectivity between large-scale networks is related to the variation in the level of suspense, the Pearson correlation *ρ* was calculated between the measured level of suspense (Fig. 7) and the connectivity time series (Fig. 8). The scan number 29 was selected as the starting state for the correlation computation, and the calculation was performed on the medium- and high-suspense level section. The correlation values are reported in Table 2.

**Table 2:** Correlation values between the suspense level and the series of weights within and between networks.

| Networks | $\rho$ value with suspense |
| --- | --- |
| Salience & Salience | 0.09 |
| Fronto-Parietal & Fronto-Parietal | 0.15 |
| Salience & Fronto-Parietal | 0.44 |
| Default & Default | -0.29 |
| Salience & Default | -0.55 |
| Fronto-Parietal & Default | -0.29 |

As observed, there is a positive correlation between suspense and connectivity weights between the fronto-parietal network and salience network, as well as within the networks. In contrast, there is a negative correlation between the connectivity weights within the default network and between the default and the other two networks.

Remarkably, when the states were associated with their communities, the intervals within the colored groups exemplified the correlation between the time series of suspense and functional connectivity. For instance, the level of suspense decreased in the lavender interval (52-72), which coincides with the tendency of connectivity weights between the fronto-parietal network and salience network, as well as within the networks, thus, they have a positive correlation.

Meanwhile, its tendency is the opposite for the connections involving the default network, showing their anti-correlation. In the same way, in the green interval, the suspense steadily increases, followed by a decrease in the blue interval, contrasting with the behavior presented by the connections between default and the other two networks, thus, an anti-correlated tendency.

Furthermore, when we used a standard change point detection algorithm [30], we retrieved six transitions where four of them coincide with bridge states given by the communities.

For instance, the transition between yellow and orange communities spots the change from low-level suspense to middle and high suspense. Importantly, the all-time series present a similar trend in this part of the scan.

### Validation against null models

A statistical test validated that the reported results were due to real changes in the FC of the brain. In this test, the null hypothesis corresponded to the static FC; meanwhile, the alternative hypothesis corresponded to the FC being dynamic [22]. For approximating a suitable null distribution, a large number of phase-randomized surrogate fMRI data was generated following the method introduced in [31] where all linear auto-correlations and cross-correlations time series were preserved, adding the same phase perturbations to all regions’ time courses for each participant.

The full pipeline of analysis (Fig 1**C-G**) was applied to the surrogate fMRI data simulating the null distribution. In particular, when generating the Mapper graphs, parameters *n* and *o* were not tuned but multiple combinations of the parameters were considered. In Fig 9 we present some Mapper graphs generated by the surrogate data.

**Figure 9.**
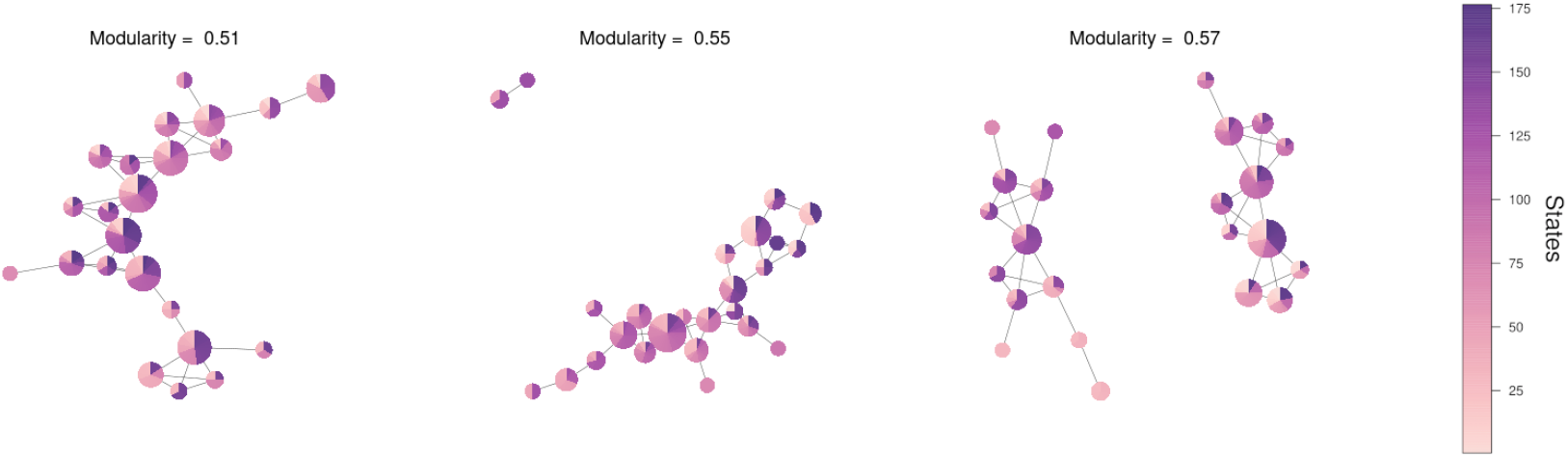
Some Mapper graphs generated by the surrogate data approximating the null model.

Results suggest that these null data (1) did not demonstrate significant variations in FC, (2) did not show a correlation between the dFC between large-scale networks with the level of suspense, and (3) did not present a communal structure in the graph’s outputs of Mapper. Thus, (1) the variance present in the functional connectivity time series for each pair of regions in the networks, (2) the correlation values between the variation of suspense and changes in the connectivity between pair of regions, and (3) the modularity of the communities in the graph (Fig 5) are all statistically significant (*p <* 0.05).

## 3 Discussion

Inspired by the psychological model of suspense by Lehne and Koelsch [1] and its consistency with a network-based approach of the emotional brain, this study investigated the hypothesis that the emotional experience of suspense depends on dynamic interactions within and between large-scale networks. For this purpose, Mapper, a topological-based tool capable of encapsulating data’s topology in a graph, was used to represent the dynamic across time of the FC. This process resulted in a graph representing the structure of the temporal dynamic of the FC. Then, community structures characterizing possible regimes in the FC dynamic were determined. Finally, changes between large-scale networks involved in the emotional expereince process and the level of suspense were related to these topological features.

### Brain dynamics are related with suspense

A central finding of our study is that the functional connectivity within and between salience, fronto-parietal and default networks changed dynamically as the level of suspense varied. Identifiable results are found in the works of McMenamin et al. [11] and Najafi et al. [12] in unraveling the dFC of the same large-scale networks during anxious anticipation using an instructed threat of shock paradigm. Similarly, our results are in agreement with the brain activation states during suspenseful film viewing found by Bezdek et. al. [5].

The connections between the salience and fronto-parietal networks have a positive correlation with the level of suspense, indicating the communication between the two networks increased in the presence of the stimulus of suspense. Meanwhile, the connections’ weights between the default network with the salience and fronto-parietal networks had a large and moderate anti-correlation with the level of suspense, respectively. Thus, the increasing suspense is associated with the segregation of the default network from the salience and fronto-parietal network, at the same time, the decrease of the suspense prompts again the communication between the networks. Both results are in line with Najafi’s work.

Conversely, the functional connections within the salience network have a small correlation value with the suspense’s level but have a sustained level connection during suspense. This result is not in line with the one reported by Najafi et. al. where the connections within the network presented the strongest change, increasing in periods of anxious anticipation and decreasing when the threat creating anxiety retreated. However, it agrees with McMenamin’s finding were threat did not have consistent effects on the salience network.. A potential reason is that the movie stimulus is motivationally salient, peaks of suspense signaled potential dangerous events, meanwhile, troughs of suspense signaled safety and hopeful events. Notably, this evidence aligns with the results reported by Bezdek et. al., where peaks of suspense are associated with brain states marked by high activity in the salience network.

In the same way, the correlation of the suspense with the connections within the default network is negative, which corresponds to with the result in [5] where the deactivation of the default network occurred more frequently at peaks of suspense meanwhile its activation was more common at valleys of suspense. This suggests the communication within the default network decreases as the level suspense increases. In fact, the regions in the default network have connectivity weights in the interval between 0.2 and 0.3. These values are low in comparison to the other series. The thresholding is partially responsible for the low values, since most of the values that were reduced to 0 belong to the default network, then the average values of connections diminished, however, this fact indicates that the lower values on the matrices are precisely in the default network. We consider that it indicates that the communication within and between the default network during the scan is not strong and could be related to the low activity of the default network during externally oriented tasks [32]. In particular, the regions in the temporal lobe have the lowest connectivity values, especially with the salience and fronto-parietal regions, which it is consistent with the fact that temporal regions are not usually consider inside the default network given the small connection with the other regions within the network.

### Brain dynamics shape the Mapper graph

Remarkably, the communities found in the Mapper graph are completely characterized by the differentiated behavior of the connections between large-scale networks (Fig 6) and the level of suspense. Hence, the presence of communities could represent six different phases of FC states the brain undergoes when experiencing suspense, and transition states are in reality connectivity configurations that the brain goes through between more stable phases. Indeed, most of the transition states indicate moments in which both the connections between large-scale networks and the suspense have abrupt shifts. Unexpectedly, the blue and solid green communities do not share any state even though they are contiguous in time. Separation between communities is especially associated with a steep rising slope in the connectivity weight between salience & default and fronto-parietal & default networks. We conjecture that the change is so fast that our window width is not able to capture the transition state between them.

Besides, some communities comprise states from different intervals of time, thus, we expect these intervals to have the same characteristics. For example, both blue intervals start with a decreasing level of suspense; meanwhile, the connections’ weights increase and decrease accordingly to their correlation with suspense. We hypothesize that the brain can revisit phases and it is possible to determine the stimuli associated with these shifts.

Importantly, previous works in suspense processing also describe the existence of brain states emerging during the processing of suspense stimuli [5]. In this case, these brain states correspond to clusters explaining the instantaneous whole brain volume activity registered using a high temporal and spatial fMRI sequence. In contrast, the brain states described for the proposed analysis aim to describe the dynamic functional connectivity of brain region interactions over time, not the brain activation itself.

Finally, given this behavior of the groups by time intervals, we computed a clustering directly from our distance matrix (Table 1) finding an almost bijection between the intervals from the clusters and those from the communities. Having said that, some studies have used this hierarchical clustering approach [33, 34] or k-means [35, 36] to summarize the states obtained into smaller sets. Hence, the Mapper approach has the same ability as the usual clustering methods to recover the geometrical structure obtained from the metric space. More significantly, Mapper does not constraint the number of sets and it captures the transitions between phases. Simply identifying clusters may not be sufficient to determine the connections and shifts between them.

In this way, Mapper generates a combinatorial object (graph) that reveals the presence of an organized dynamic transition between phases of FC states, which can be characterized by a differentiated behavior of the FC between and within large-scale networks related with the change in the level of suspense. Moreover, the graph summarizes the temporal evolution of the brain through these phases and highlight the key states present in the transitions between them.

### Brain dynamics and TDA

Recently, TDA was explored to characterize fMRI brain dynamic in resting and during stimuli regimes [17–20, 37–41]. As expected, these different analyses focused on particular dynamical aspects of the fMRI dynamic. For instance, recent works describe the temporal dynamic underlying the instantaneous whole-brain dynamic, i.e., the activation observed in the whole brain at particular TRs [17, 19, 20]. Our work also aims to define a temporal dynamic but for the interactions among regions from networks previously known to be involved in suspense processing. Therefore, we focused on the short-time FC dynamic over time exhibited during emotional processing.

Other TDA-based works aimed to describe the topology underlying the relationships between different brain areas, both at an instantaneous level [40] or by considering the complete temporal dynamics exhibited by each region [38, 41–43]. Because our primary interest was to describe the temporal dimension of the suspense processing, the proposed TDA analysis focused on time and not on regions. However, future work may consider a higher-order description of the FC computed on the slicing window.

Finally, other recent TDA-based works also focused on describing FC’s temporal dynamic over time [37, 39, 44]. Nevertheless, these works did not account for the possible relationship between stimuli and fMRI FC dynamic, as they focused on describing the resting-state activity [37, 39], or did not consider high-order relationships provided by graphs [44], which in the proposed analysis provided information not only of the groups of states but also about possible transitions among them.

### Limitations and future directions

The results presented here must be considered in the context of experimental and methodological limitations. To begin with, the movie we used as stimuli was originally designed to evoke suspense [45], but not to contrast levels of suspense, in contrast to [5] where the experiment was designed to have defined periods of peaks and valleys of suspense. In the movie, we had a short period of low suspense followed by a period of sustained high suspense; thus, we had a transition from low to high suspense, but not the opposite. Besides, in the second part we have difficulty in identifying the transitions between peaks and troughs of the level suspense. This fact in combination with the low temporal resolution of fMRI poses an upper limit for our ability to resolve the neural effects of suspense in time. Our results suggest future research to design experiments with multiple blocks of low- and high-suspense, including periods of transition from both states.

Furthermore, our results are circumscribed to Shen Atlas, used to define brain regions and the selection of brain regions for each network based on the work of Power et al. [46]. However, we hypothesize the results will be similar for a different selection of atlas based on functional connectivity and network separation since we did not consider roles for particular regions but general behaviors of connected regions. Nonetheless, an interesting path for future research could be considering the hypothesis of overlapping networks, where specific regions belong to several intersecting networks and its participation at a given time is context dependant [47].

Additionally, future research should investigate the contributions of subcortical regions linked to emotional experiences [6] such as the amygdala, PAG, BNST, hypothalamus, and thalamus in dynamic interactions of networks in the presence of suspense. In addition, the work could be extended to recognize the contribution of the regions inside each network, e.g., the Frontal Orbital Cortex inside the salience network.

Another venue for future research is the generation of Mapper graphs that encapsulate the brain dynamics for only one participant at a time and analyze how individual differences influence the topological properties of the graphs. For example, considering the mental health of the participants. Similarly, an average Mapper graph can be created for age-binned groups and capture age-related variability in the brain dynamics.

Finally, the proposed methodology could be apply in the research to the study of the dynamic processes related to emotions like anger, happiness, and sadness, among others. Looking forward, it can be argued, we could characterize each emotion by the temporal transitions between different connectivity states [48].

### In conclusion

Our TDA-based approach revealed a dynamical organization of the brain across the emotional experience of suspense, specifically, the changes in the functional connectivity within and between the salience, fronto-parietal, and default networks are correlated with of the level of suspense. In this way, our study supports the idea of understanding emotional experiences as dynamic interactions of domain-general neural systems and highlights topological-based methods as powerful tools to visualize and recognize the dynamic brain networks.

## 4 Material and methods

### Data acquisition

#### Participants

We analyzed an open dataset acquired by the Cambridge Center for Aging and Neuroscience (Cam-CAN) [23, 24]. A total number of 700 participants were selected to take part of this study including 100 individuals in each decile age 18-87 and an equal number of men and women. Eligibility measures included cognitive health, meeting hearing and English language requirements, and being eligible for MRI scanning [24]. The final sample for the fMRI time series analysis included 492 participants.

#### Stimuli

The stimulus is an edited version of Alfred Hitchcock’s “Bang! You’re Dead”, a black and white television drama portraying a little boy that plays around with a charged revolver thinking it is a toy gun (Fig. 2**A**). The entire 25 minute episode was condensed to 8 minutes while maintaining the narrative. Participants were instructed to watch, listen, and pay attention to the movie (they were not aware of its title) [23].

#### Continuous Response Measurement Sample

A total of 22 human raters were recruited to provide continuous response measures [49]. When viewing the same edited clip from the fMRI sample, participants were directed to “continuously evaluate the degree of suspense “they were experiencing using an online rating tool [50]. We use the averaged degree of reported suspense calculated in [4]. (Fig. 2**B**)

#### fMRI scanning and processing

MRI data were acquired using a 3T Siemens TIM Trio System with a 32-channel head coil. For the functional scan, T2*-weighted echo planar images (EPIs) were acquired using a multi-echo sequence (TR = 2.47 seconds; 5 echoes; flip angle 78 deg; 32 axial slices; field of view = 192 mm × 192 mm; voxel-size = 3 × mm 3 mm × 4.44 mm) with an acquisition time of 8 minutes and 13 seconds. More details are in the report on [23].

The imaging data were processed using the nipype framework [51] including motion correction, slice-time correlation, co-registration and nonlinear normalization to the MNI template. Functional data were detrended and high-pass filtered at 0.01 Hz and time series were extracted using NiLearn [52]. From the original dataset of 646 participants some participants were excluded if the data were incomplete or exhibited abnormal behavior, resulting in a final sample of 492 participants with 193 volumes of 231.502 voxels. We use the processed data from [4].

### Regions of interest

The whole volume is divided in 268 regions the Shen et al. atlas [53], divided into nine networks [46]. Information of each region is retrieved from the BioImage Suite Web [54]. We focused on salience, fronto-parietal and default networks. We excluded some regions that are not commonly linked to these networks in the literature. Nevertheless, we maintained the temporal regions in the fronto-parietal and default networks. Ultimately, we considered 74 regions from the Shen Atlas (Table 3)

**Table 3:** Regions considered for each network. If R or L are not specified, then both hemispheres are considered.

| Salience |  |  |  |
| --- | --- | --- | --- |
| 1 | Angular Gyrus R | 7 | Frontal Orbital Cortex |
| 2 | Paracingulate Gyrus | 8 | Dorsolateral superior Frontal Gyrus |
| 3 | Inferior Frontal Gyrus | 9 | Medial superior Frontal Gyrus L |
| 4 | Middle Frontal Gyrus | 10 | Cingulate Gyrus ant. Division R |
| 5 | Insular Cortex | 11 | Supramarginal Gyrus post. division R |
| 6 | Frontal Pole | 12 | Frontal Operculum Cortex |
| Fronto-Parietal |  |  |  |
| 1 | Frontal Pole | 6 | Inferior Frontal Gyrus pars opercularis |
| 2 | Superior Frontal Gyrus | 7 | Inferior Frontal Gyrus pars triangularis |
| 3 | Middle Frontal Gyrus | 8 | Middle Temporal Gyrus R |
| 4 | Supramarginal Gyrus | 9 | Inferior Temporal Gyrus R |
| 5 | Precentral Gyrus | 10 | Superior Parietal Lobule |
| Default |  |  |  |
| 1 | Hippocampus | 12 | Orbital superior Frontal Gyrus |
| 2 | Frontal Pole | 13 | Inferior Frontal Gyrus pars triangularis |
| 3 | Temporal Pole | 14 | Dorsolateral superior Frontal Gyrus |
| 4 | Precuneus | 15 | Medial superior Frontal Gyrus |
| 5 | Frontal Orbital Cortex | 16 | Superior Temporal Gyrus ant. division |
| 6 | Frontal Medial Cortex | 17 | Middle Temporal Gyrus post. division |
| 7 | Angular Gyrus L | 18 | Middle Temporal Gyrus ant. Division |
| 8 | Paracingulate Gyrus | 19 | Superior Temporal Gyrus post. division |
| 9 | Fusiform gyrus | 20 | Cingulate Gyrus post. division |
| 10 | Lingual Gyrus | 21 | Parahippocampal Gyrus post. division |
| 11 | Middle Frontal Gyrus |  |  |

### Forming connectivity matrices

The final dataset consists on *P* = 492 patients, each with a scan of *N* = 74 regions tracked in *T* = 193 volumes. Thus one participant is represented by an *N* × *T* matrix, that is, regional activity magnitudes as a function of time.

#### Sliding window method

To asses FC between regions we used Pearson correlation, a statistic measure of linear correlation between two sets of data. Given two samples *X* = {*x*_1_,…, *x*_*n*_} and *Y* = {*y*_1_,…, *y*_*n*_} the Pearson correlation between *X* and *Y* is defined as

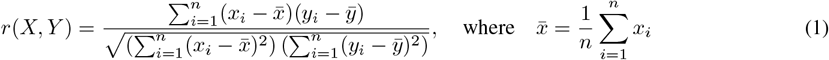

Now, in the framework of the SW, we select a window of size *W*. Then, we define a *N ×W* matrix representing the regional activity within the temporal span from time *t* = 1 to time *t* = *W*. Subsequently, we compute the FC between each pair of regions. Afterwards, the window is shifted (slided) by a step *S*, and the same calculations are repeated over the time interval [1 + *S, W* + *S*]. This process is iterated until the window spans the end part of the time series. This procedure yields *U* = *T*− *W* + 1 windows each with *N*× (*N* −1)*/*2 values, which are summarized into a connectivity matrix describing the state of the brain during the examined temporal interval.

For the parameters *S* and *W* we chose *S* = 1 because this step has been proved to be the optimal value to detect connectivity changes between the windows [55]. On the other hand, *W* is a matter of debate, since the output of the SW can be highly dependent on it. Too short window lengths increase the risk of introducing spurious fluctuations [56], in contrast, too long windows are not capable to capture short-lived FC variations. Nevertheless, previous studies suggested that window sizes around 30 − 60 s are able to produce robust results in dFC and in most cases, different window lengths, when chosen in this interval stabilize and do not yield substantially different results [14, 57]. Consequently, we choose *W* = 18 volumes which gives an span of approximately 44, 5s falling into the recommended interval as we wanted. Hence, we generated *U* = 193 − 18 + 1 = 176 FC states.

#### Average over all participants

Afterwards, we used the SW over all individuals. Since the correlation matrices among the patients may be quite variable, to smooth and stabilize the correlation coefficients between the regions [58], we take the average of all connectivity matrices for each state. For this average, we first applied Fisher’s transformation to all correlations, then we average and subsequently back-transform the values. In this way, we reduce the bias induced by the bounds of correlation at −1 and 1 [59].

#### Thresholding connectivity matrices

Having estimated the connectivity matrices for each state, we need to apply a global *threshold τ* in order to dismiss negative connections and spurious or weak positive connections. Let *C* = [*c*_*ij*_] be a connectivity matrix. We form the matrix *Ĉ* = [*ĉ*_*ij*_]

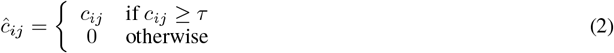

Now, with the purpose of choosing a proper *τ* we consider the notion of connection density. We can define the connection density *κ* as

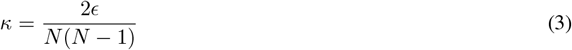

where *ϵ* is the total number of weights which weight is equal or greater than *τ*.

From [60] we know that brain connectomes typically have connection densities below 0.5 since over this value they present random topologies. Given that the density increases when *τ* increases, we choose the minimum *τ* such that the density of all states does not exceed 0.5. Specifically, we start at *τ* = 0.3 and increase *τ* at a step of 0.1 until every state’s density is below *κ* = 0.5.

Nevertheless, the analysis of the dataset without the threshold revealed the same topological shape (see next subsection), confirming the threshold process removed noisy connections without losing any meaningful information of the data.

### Revealing the shape of data using topology

The data set consist on 176 connectivity matrices of size 74 ×74 pair of regions. As a initial step, we calculated the dissimilarity between the matrices in form of vectors using the Manhattan distance. Then, we formed a distance matrix with 176*/*2 × 175 = 15400 entries.

#### Persistent homology

Seeking to extract the shape of the data set we compute its homology: an invariant property of topological spaces that permit us to recognize its shape. Informally, for a space *X, H*_*i*_(*X*), a homological class, is the vector space associated with the *k −*dimensional holes of *X*. In particular, the dimension of *H*_0_(*X*) is the number of path connected components of *X*, the dimension of *H*_1_(*X*) is the number of circles, the dimension of *H*_2_(*X*) is the number of spheres and so on.

To extract the homology of a finite data set we work with simplicial complexes. A *simplex* of dimension *d* is a set of *d* + 1 elements. A *simplicial complex S* is a finite collection of simplices such that if *α* ∈ *S* and *β* ⊂*α* implies *β* ∈ *S*. The dimension of the complex is the maximum dimension of any of its simplices. For example *S* = [*v*_0_, *v*_1_] = {{*v*_0_}, {*v*_1_}, {*v*_0_, *v*_1_}} is a simplicial complex of dimension 1 or a 1-simplex. Simplicial complexes can be visualized as d-dimensional polytopes: [*v*_0_] as a point, [*v*_0_, *v*_1_] as a line segment, [*v*_0_, *v*_1_, *v*_2_] as a triangle, [*v*_0_, *v*_1_, *v*_2_, *v*_3_] as a tetrahedron, and so forth. In a simplicial complex, we can consider the *k −*dimensional holes as voids bounded by simplices of dimension *k*. In dimension 0, are components connected by segments, in dimension 1, are cycles bounded by segments, in dimension 2, are voids bounded by triangles and so on.

Finally, to associate a simplicial complex to a finite set we choose a real number *ϵ*, we create a covering generating an open ball of radius *ϵ* centered in each data point and then we add the simplex [*v*_0_, *v*_1_,…, *v*_*i*_] if the balls of the data points *v*_0_, *v*_1_,…, *v*_*i*_ have a not empty intersection. Subsequently, we compute the homology of the simplicial complex corresponding to *ϵ*. We repeat this algorithm for 0 *< ϵ < R* for some *R*. Then, the homological classes that *persist* trough a large range of *ϵ* reveal the homology of the data set.

#### Mapper construction

After forming the metric space the second Mapper step is filtering. We used Isometric Mapping (Isomap) for dimensional reduction. The method is a combination of calculating the geodesic distance between data points and then projecting them to 2D space using multidimensional scaling. In particular, the input of the algorithm is the distance matrix calculated before and the geodesic distance used *k* = 30 as the number of nearest neighbors which highlighted the homological properties found using persistent homology.

Subsequently, the filter space is divided into overlapping bins that cover the collection of points (Fig. 1 **E**). The covering possesses two parameters: the number of intervals (*n*) that each dimension of the parameter space is divided in and the percentage of overlapping between the intervals called overlap (*o*). We tuned these parameters so that we preserved the shape of the data derived from *persistent homology*. Thus, we generated 25 different covers of the parameter space varying the intervals from 4 to 8 (steps of 1) and the overlap from 25 to 45 (steps of 5). Afterwards, we localized the Mapper graphs that resemble the objective homology with the minimum amount of nodes. This approach selected the parameters *n* = 6 and *o* = 35.

The fourth step is applying partial clustering within each bin to recover the geometrical information missed after filtering but yet reducing the complexity of the data. We applied the agglomerative hierarchical clustering. In this method (1) each data point as his own cluster, (2) clusters get merged if they have the minimum distance between all clusters, (3) we calculate the distance between this new cluster and the rest and (4) we repeat this process until we have only one cluster. There are different types of methods depending on how we calculate the distance between two clusters. For example, in the single-link method [61], the distance between any two clusters *C* and *D* is defined as the minimum distance between the elements in *C* and *D*:

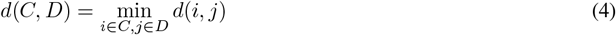

In contrast, the complete-link method defines the distance between clusters using the maximum.

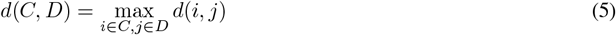

Note that in step 2 for each merging distance we have a different set of clusters. To chose the proper division of the data we have used the cut-off method proposed in the Mapper implementation called *continuous*, where we have estimated the density distribution of the merging distances and chose the one with the smallest value.

Finally, the clusters form a covering of the data set, thus, we can associate a simplicial complex to it. Since we are only using 0- and 1-simplices, the topological object has the shape of a graph where each cluster is a node and they share an edge if they share data points

#### Software

For this construction, we use the implementation of Mapper developed by Piekenbrock et. al in R language [28, 62]. Besides, for the filter we use the implementation of Isomap found in the Python’s package scikit-learn [63] and Python’s package ripser [64] to calculate the persistent homology.

### Exhibiting geometric organization

To find the communities in the graph,we have used an approach of random walks [29]. Intuitively, random walks on the graph tend to get “trapped” in communities. First, a discrete diffusion process is applied to the graph to define the random walks. Then, we can determine the probabilities of going from node *i* to *j* and from *j* to *i* through a random walk. This probability is used to define an efficiently computable distance between nodes *i* to *j*. Subsequently, the distance is used in a hierarchical clustering algorithm (merging nodes to communities); thus, each merge defines different communities. Finally, the partition that maximizes modularity *Q* is chosen.

Modularity is a well-known measure to determine the quality of the partition given by the communities [65]. *Q* is defined as

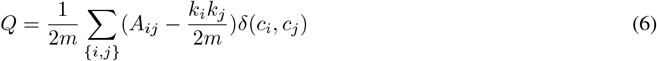

where *m* is the number of edges, *A*_*ij*_ is the adjacency matrix where *A*_*ij*_ = 1 if {*i, j}* is an edge an 0 otherwise. *k*_*i*_ is the degree of *i, c*_*i*_ is the component *i* belongs to and *δ*(*x, y*) = 1 if *x* = *y* and 0 otherwise.

Finally, community detection and modularity values were calculated with the igraph package [66] for *R*.

### Relating the level of suspense to the shape graph

To connect the level of suspense and the communities found on the Mapper graph we detect the change points on the time series of level of suspense and related with the limits of the time blocks defined by the communities. To detect such change points we use algorithm based on a pruning rule which minimizes a piecewise linear model with unknown number of changes. We use an implementation found in ruptures [30], a Python based package.

### Linking the shape graph to dynamic functional connectivity

To calculate the FC within a network *N*_1_ or between networks *N*_1_ and *N*_2_ we use connectivity weights defined in [12] as follows:

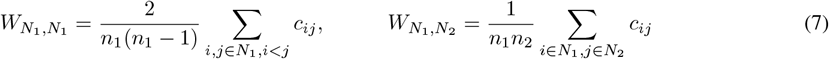

where *n*_1_ and *n*_2_ are the number of regions in *N*_1_ and *N*_2_ respectively and *c*_*ij*_ is the measure of functional connectivity between regions *i* and *j*.

To calculate the dFC within and between networks we measure their connectivity weights trough time calculating 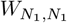 or 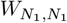 for each state along time.

### Validation against null models

To validate that our results are due to real changes in the FC of the brain we define a “statistical hypothesis test in which the null hypothesis corresponds to the correlation being static and the alternative hypothesis corresponds to the correlation being dynamic” [22]. To start a proper null distribution is needed, one that corresponds to the static FC. Such null distribution is approximated by surrogate data. To generate it, for each patient, all time series corresponding to each brain region are phase-randomized by the same phase. We repeat this process on all 492 patients. Then, we iterate this process a thousand times.

The time series were phase-randomized following the method introduced in [31]. In this method, given a time series *x*(*t*) we take its discrete Fourier transform, then we we rotate the phase *ϕ* at each frequency *f* by an independent random variable *ψ* which is chosen uniformly in the range [0, 2*π*) and we obtain the surrogate time series 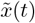 calculating the inverse Fourier transform

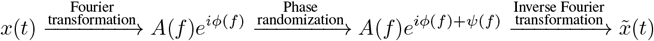

By an extension of the Weiner-Khintchine theorem to preserve all linear auto-correlations and cross-correlations we only need to add the same phase *ψ*(*f*) for all regions for each subject.

The second step is estimating different measures of the null distribution..

1. The dFC of the surrogate data is measured using *variance η*, since it is the most widely used and the most straightforward [22].
2. The correlations of the dFC with suspense is measured after the sliding window method (Fig 1**C**) was applied to the surrogate data.
3. The modularity of the communities found in the Mappers graph generated by the surrogate data is measured. For that, the pipeline of analysis (Fig 1**D-G**) was applied. In particular, when generating the Mapper graphs, parameters *n* and *o* were not tuned but multiple combinations of the parameters were considered.

The third step is deciding if each measure of the results is statistically significant or not. For instance, let *η*_0_ be the variance of the data. If *η*_0_ falls within the 5% highest values or lowest values (if the test is one-sided) of the variance distribution of the null model we can conclude it is statistically significant [22]. Otherwise, it is not. As usual we note the significance level as *p* (percentile) and we choose 0.05 as our threshold. In this way, we reject *η*_0_ if *p >* 0.05.

## Availability of data and materials

The raw fMRI files are available in the Cam-CAN repository https://www.mrc-cbu.cam.ac.uk/datasets/camcan [24].

The processed data and the continuous response measurement are available on request from the corresponding authors [4].

Code to reproduce and document the analyses is accessible online at https://github.com/aaolaveh/TDA_suspense.

